# Strategies for resolving cellular phylogenies from sequential lineage tracing data

**DOI:** 10.1101/2025.01.14.632902

**Authors:** Nicola Mulberry, Tanja Stadler

## Abstract

A combination of recent advancements in molecular recording devices and sequencing technologies has made it possible to generate lineage tracing data on the order of thousands of cells. Dynamic lineage recorders are able to generate random, heritable mutations which accumulate continuously on the timescale of developmental processes; this genetic information is then recovered using single-cell RNA sequencing. These data have the potential to hold rich phylogenetic information due to the irreversible nature of the editing process, a key feature of the employed CRISPR-based systems that deviates from traditional assumptions about molecular mutation processes. Recent technologies have furthermore made it possible for mutations to be acquired sequentially. Understanding the information content of these recorders remains an open area of investigation. Here, we model a sequentially-edited recording system and analyse the experimental conditions over which exact phylogenetic reconstruction occurs with high probability. We find, using simulation and theory, explicit parameter regimes over which simple and efficient distance-based reconstruction methods can accurately resolve the cellular phylogeny. We furthermore illustrate how our theoretical results could be used to help inform experimental design.

## 1. Introduction

Lineage tracing tools allow researchers to track the development of organisms and tissues at the resolution of single cells. Uncovering the underlying single-cell phylogeny is key to understanding many complex cellular dynamics, as the tree provides a robust statistical framework in which to analyse traits which emerge and change as the cell population grows and differentiates (Stadler et al., 2021). While the phylogeny can be traced under the microscope in simple settings of few cells and short timescales (McKenna and Gagnon, 2019), researchers must turn to other data sources to re-construct phylogenies in more general settings. Naturally occurring genetic variation within a single cell typically accumulates over timescales much longer than a typical cell’s life-cycle. To investigate developmental processes in depth, researchers can instead use genetic engineering techniques to equip cells with artificial “barcodes” for the explicit purpose of capturing the lineage. These technologies have the potential to produce data with strong phylogenetic signal, and mathematical modelling can be used to analyse the experimental conditions under which accurate reconstruction of the underlying tree is possible.

Dynamic lineage tracing technologies leverage the ability of CRISPR-based systems to induce heritable mutations (‘edits’) into compatible locations in a genome (McKenna and Gagnon, 2019; Chen et al., 2022). We will refer to each of these loci as a ‘tape’, and the combination of all recording loci (tapes) together with the editing machinery in a cell will be referred to as the ‘recorder’ or ‘recording system’. Once a target site on a tape is edited, it is no longer recognizable by the editing machinery. In this way, edits accumulate irreversibly. Recent technological advancements furthermore capture the order of editing events (Choi et al., 2022; Loveless et al., 2021), thereby increasing the phylogenetic information content of the system.

Despite the irreversible and sequential nature of edits induced by the systems developed in Choi et al. (2022) and Loveless et al. (2021), exact phylogenetic reconstruction from these data remains a challenge. In particular, the system of Choi et al. (2022) is characterized by a limited number of target sites per recorder. This requires careful calibration of editing rates in order to ensure sufficient diversity is accumulated for the duration of the entire experiment. An editing rate that is too low may generate insufficient information to resolve phylogenetic relationships, while an editing rate that is too high leads to tapes which become saturated before the end of the experiment. Another key feature of these systems is the diversity of the induced mutations. It is challenging to infer phylogenies from tapes with a low diversity of mutations, due to the increased potential for sites sharing identical edits through chance rather than through inheritance (a process generally referred to as homoplasy). Low diversity of mutations may happen if the space of possible edits is small, if certain edits are inserted much more frequently than others, or through a combination of these two factors. Although the sequential nature of edits makes identification of homoplastic sites possible in certain circumstances, the possibility of homoplasy occurring and interfering with the correct identification of a cell’s lineage cannot fully be ignored. All of these features, among others, can be explored using an appropriate model of tape evolution.

The characterization of accurate or exact tree reconstruction under classical models of sequence evolution has been previously studied (for example, Fischer and Steel (2009); Dornburg et al. (2019); Townsend and Lopez-Giraldez (2010)). However, the evolution of CRISPR recorders violates many assumptions of these traditional models, and so new models and analyses developed specifically for these recording systems must be developed. Two recent papers (Wang et al., 2023; Salvador-Martínez et al., 2019) have examined the accuracy of trees reconstructed from such lineage tracing data. In particular, Wang et al. (2023) develop a model for recorder evolution and derive theoretical bounds for tree reconstruction. In this model, the authors capture the irreversibility of induced edits, but they do not consider sequential editing. Additionally, these results have limited practicality as they often greatly overestimate the amount of information necessary for reliable reconstruction of the tree topology. Salvador-Martínez et al. (2019), on the other hand, use a simulation-based approach only, and again do not consider sequential editing systems.

In this work, our goal is to assess the information content in data from sequentially edited CRISPR systems by first developing an appropriate model of sequence evolution, and subsequently deriving theoretical bounds on the reconstruction accuracy of data generated under a given set of experimental parameters. We show using simulations that this theory provides practical estimates on the reconstruction accuracy; we furthermore show how these theoretical results can be used to design and guide experiments in a way that simulations alone cannot. Our model focuses on key experimental parameters such as the number of target sites per tape, tape copy number and editing rate. We will generally assume that the number of target sites is very limited, while the editing rate can be flexibly tuned. In comparing our theoretical results to simulations, we often use the distance-based method UPGMA (Sokal and Michener, 1958) to reconstruct the tree topology. We focus on UPGMA here due to its scalability and popularity in the single-cell literature; our goal is not to assess the accuracy of any given reconstruction method, rather to show that we can find experimental conditions wherein we expect any appropriate method to be able to accurately reconstruct the tree.

## 2. Model

In dynamic lineage tracing experiments, a progenitor cell is equipped with a recorder system consisting of one or more tapes, which are passed down and accumulate edits as the cells divide, and thereby act to record the lineage. After some fixed amount of time, a sample of extant cells are sequenced. Our goal is to exactly reconstruct the cellular phylogeny from the sequences recorded on each tape. We will model a sequential editing process, inspired by the system described in Choi et al. (2022): Each tape has a fixed and finite set of target sites which are initially unedited and all but the first of which are deactivated. Edits occur randomly throughout the process, and a site being “active” means that it can be edited. Each edit subsequently activates the next site in the tape (if such a site exists). In this way, edits accumulate in sequence. Each edit is drawn from a finite and pre-defined set of possible characters.

We start with a cell equipped with *m* tapes each having *k* target sites. Initially, each site is in an unedited (“0”) state. For each tape, we assume that mutations arise at constant rate *λ* independently of when the last mutation event took place and independent of cell division events. In a given tape, let *k*^′^ be the first unedited site. During a mutation event, the character at site *k*^′^ changes from 0 to *i* with probability *ξ*_*i*_, *i* = 1, …, *j*. If no such site exists, then the tape is saturated and no mutation occurs. Throughout, we denote by *q* the probability that two independent editing events will insert the same character,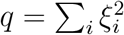. The model parameters are outlined in Table 1.

**Table 1.**
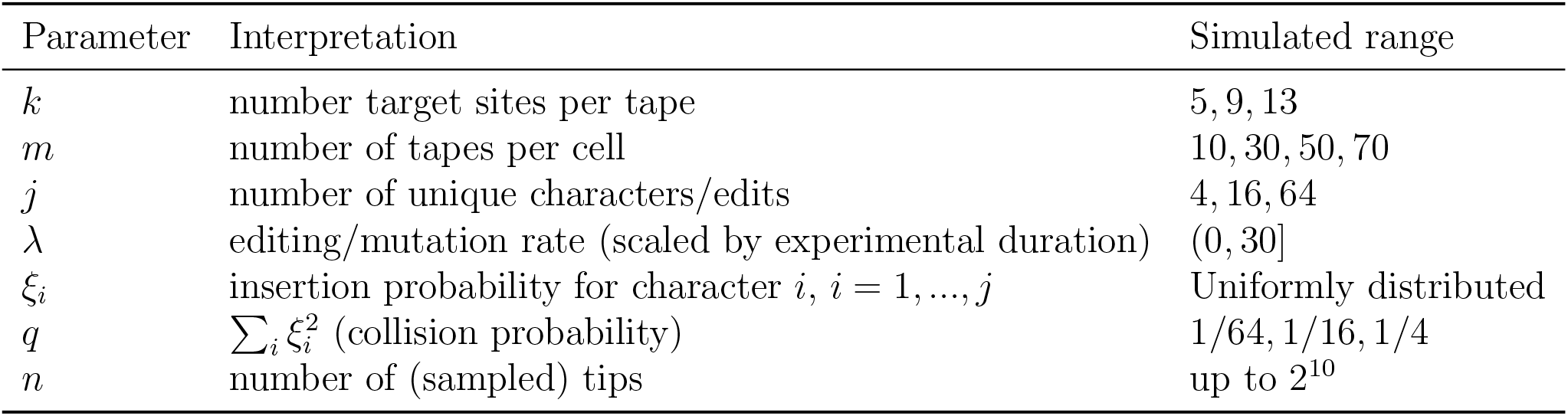
Parameters for our model of a sequentially edited recording system, and the typical ranges over which we validate the model with simulations. Our results will depend on the quantity *q*, which is defined through the insertion probabilities *ξ*_*i*_.

| Parameter | Interpretation | Simulated range |
| --- | --- | --- |
| $k$ | number target sites per tape | 5, 9, 13 |
| $m$ | number of tapes per cell | 10, 30, 50, 70 |
| $j$ | number of unique characters/edits | 4, 16, 64 |
| $\lambda$ | editing/mutation rate (scaled by experimental duration) | (0, 30] |
| $\xi_i$ | insertion probability for character $i$, $i = 1, \dots, j$ | Uniformly distributed |
| $q$ | $\sum_i \xi_i^2$ (collision probability) | 1/64, 1/16, 1/4 |
| $n$ | number of (sampled) tips | up to $2^{10}$ |

Under these assumptions, we can model the number of edits on a given tape by a censored Poisson distribution Seidel et al. (2024). Let *f*_*k*_(*x, r*) denote the probability density function for a left-censored Poisson distribution with some rate *r* (which will be defined later) and maximum value *k*, and *f* (*x, r*) denote that for the uncensored Poisson distribution with rate *r*. That is,

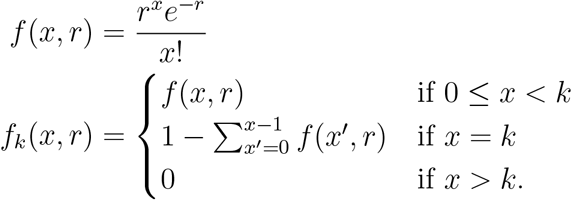

## 3. Results

We can use the model described in Section 2 to (1) derive approximate bounds on the probability of exactly reconstructing a tree in a given experimental regime, and (2) to simulate the evolution of tapes along a tree. Section 3.1 will derive our analytical results, which we validate against simulations in Section 3.2. Finally, sections 3.3 and 3.4 provide two examples of how these theoretical results could be used in practice.

### 3.1. Bounding the probability of exact reconstruction

Let 𝒯 be an ultrametric phylogeny representing a binary branching process with leaf set ℒ and internal edge set ℰ. We denote by *d*_*u*_ the depth of each node, where *d*_*a*_ = 1 for all leaf nodes *a* ∈ℒ and *d*_*o*_ = 0 is the depth of the origin, where each tape has the known state **0**^*k*^. We will define a model for the evolution of a tape along this phylogeny, which generates data at the tips. In what follows, we assume that the times between cell division events can be bounded below by parameter *ℓ*. That is, *ℓ >* 0 denotes the shortest cell cycle. Our goal is to exactly reconstruct the topology of 𝒯 from the data at the tips.

We will derive two expressions for approximating the reconstruction accuracy of this system. The first will consider the case where there is no chance of homoplasy, which simplifies our approach. We then use this analysis to motivate a more general result. We validate our results *in silico*, using UPGMA to reconstruct each tree from simulated recording systems. Our general procedure will be to first work out the probability of exactly identifying a given triplet in the tree: given the sequence data at three leaf nodes *a, b, c* ∈ ℒ, what is the probability (under the model described in Section 2) that we will be able to correctly identify which pair of nodes are more closely related? Once we have this quantity, we can bound the probability of exactly reconstructing the entire tree. Our results are not rigorous bounds *per se*; our first result is only valid in an idealised parameter regime, although we find that it tends to perform well when compared to simulations. Our second result is more general but is approximate, although we will justify these approximations using simulations.

Consider the generic triplet (*a, b*| *c*) shown in Figure 1, where (*a, b*) has least common ancestor *v*, and (*b, c*), (*a, c*) each have least common ancestor *u*. Given the irreversible nature of the editing system, the triplet will be resolved–if possible–by the edits which occur between *u* and *v* (the yellow path in Figure 1). The minimal internal branch length *ℓ* will thus be a key determinant in the reconstruction accuracy. Complicating this scenario, however, is the fact that an identical subsequence of edits could arise independently from along the path (*u, c*).

**Figure 1.**
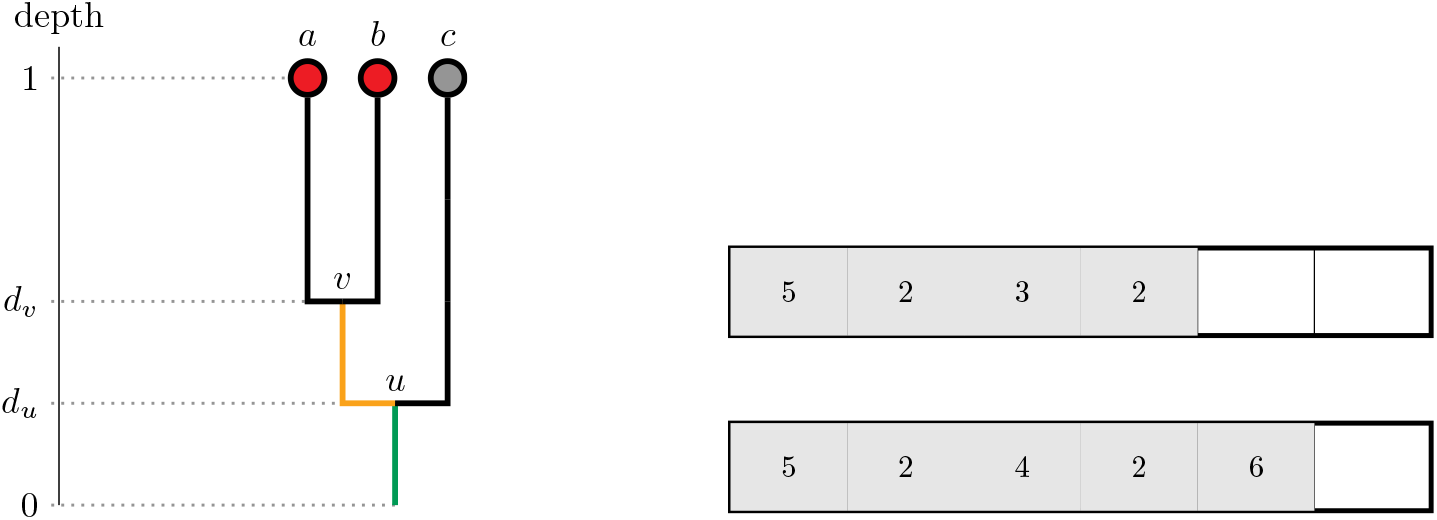
(Left) Diagram of a triplet (*a, b*| *c*). Time is always scaled so that the tips of the tree are at depth 1, and the origin of the tree (the initial cell with no edited sites) is at depth 0. Under the editing model, all edits which occur along the green path from the origin to *u* will be shared among all three nodes. All edits which occur along the yellow path from *u* to *v* will be shared between *a* and *b*. (Right) Example tapes with *k* = 6 from two different cells at the end of the experiment. The number of shared edits between the two tapes here is 2: we do not count any shared characters outside of the first shared subsequence, nor any unedited sites.

Let *D* be a distance matrix induced by the mutation process, and let 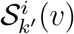denote the character at site *k*^′^ on tape *i* for any *v* ∈ ℒ. For each pair of leaves, the number of shared mutations is the length of the shared subsequence, summed over each tape and ignoring any unedited (zero) sites. That is, for any pair of leaves *a, b* ∈ℒ, we define

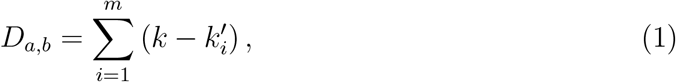

where 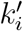is the largest index on tape *i* such that 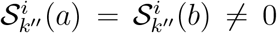for all sites 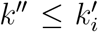. As a sufficient condition for tree reconstruction, we will require that every triplet is identifiable. That is, we will say that the tree can be exactly reconstructed if

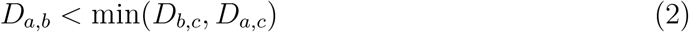

holds for all (*a, b*|*c*) ∈ 𝒯.

#### 3.3.1. Infinite characters (B_∞_)

We first consider the case of having an infinite space of possible edits, i.e. *q* = 0.

Let 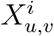 be the number of shared edits along an edge (*u, v*) ∈ ℰon the *i*th tape. Note that 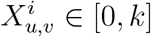for all (*u, v*) ∈ ℰ. Now consider the probability of there having been no edits along this edge. This happens if either the tape is already saturated (i.e. has accumulated *k* edits by depth *d*_*u*_), or the tape is not saturated but no editing event takes place along the path. Let *ℓ* be the minimum branch length and *d* = 1 − *ℓ* be the maximum depth of the start of each internal edge. For any edge (*u, v*) ∈ ℰ, we have

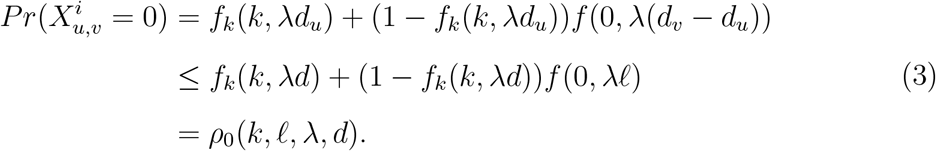

Therefore, the probability that there are no edits along a given branch on any of the *m* independent tapes is no greater than 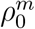. In general, we could consider *d <* 1 −*ℓ* if we do not need the entire tree to be resolved. Unless otherwise stated, we will take *d* = 1 − *ℓ* and consider the problem of resolving the entire tree. We now notice that a split in 𝒯 at depth *d*_*u*_ ∈ (0, *d*] is resolved by any edits occurring along the path (*u, v*).

As soon as at least one edit occurs on at least one tape along this path, then every triplet with least common ancestor *u* will satisfy Eq. (2) and hence be resolvable. This is due to the irreversibility of the process, and the fact that we here take *q* = 0 and so can neglect any chances of homoplasy. Hence, we can interpret 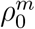as an upper bound on the probability of not resolving a split. We now state our first reconstruction bound:

##### Result

*(B*_∞_*) Given the tape mutation process described in Section 2 with q* = 0, *the probability of exactly reconstructing the phylogeny 𝒯 on n tips is at least*

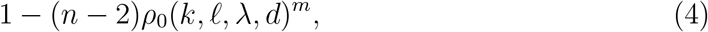

*where ρ*_0_ *is given by Equation 3*.

To bound the probability of exactly reconstructing the tree, we traverse down the tree, starting at the terminus of the stem edge, and take a union bound over the (*n*−2) internal branches. We take a union bound here since we cannot assume independence between different splits, which leads to Equation 4.

#### 3.1.2. General case (B_q_)

We now consider the more general case where 0 *< q <* 1, and so we have to take into account the chance of homoplasy occurring. We will consider the triplet (*a, b* |*c*) where *u* = *LCA*(*a, b, c*) and *v* = *LCA*(*a, b*) as shown in Figure 1. We will derive the probability that the total number of shared edits between *a, b* is less than or equal to *b, c* or *a, c*. Let 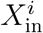denote the number of shared edits on tape *i* between *a, b* which arise after the three lineages diverge at *u*, and 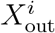 denote those between *c* and *a* (or similarly between *c* and *b*). Then 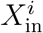 will be dominated, similarly to the case above, by the number of edits arising along the path (*u, v*). Unlike the case above, however, there may also be shared edits which arise along the paths (*v, a*) and (*v, b*) independently.

We first work out the probability of an identical sequence of edits arising independently along two paths. Let *n*_*a*_ and *n*_*c*_ denote the number of edits acquired along each path, respectively, and without loss of generality let *n*_*a*_ ≥ *n*_*c*_ ≥ *x*. Given *n*_*a*_, *n*_*c*_, the probability that there is an identical sequence of length *x* (starting from the first position) is

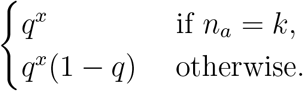

That is, the first *x* edits are identical and are followed by one different edit if the tape is not saturated (for example, see Figure 1). Given that there are maximum *k*^′^ possible edits, the probability of an identical sequence of length *x* arising on two paths of length (1 − *d*) is

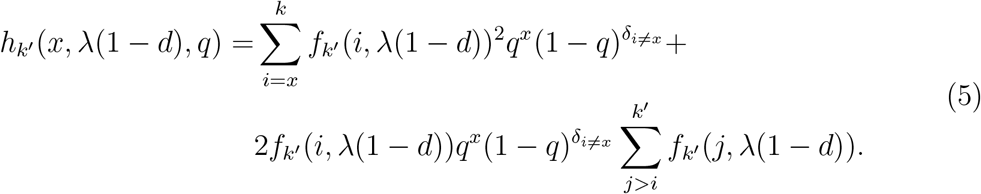

The first term looks at all cases where the two paths have equal length, and the second term looks at all cases where the second path is strictly longer than the first. In both cases, we multiply by 1 −*q* only if the length of the shorter path is strictly greater than *x*.

To find 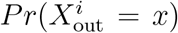, we see that any shared sequence which arises between *a* and *c* does so independently along the paths from *u* to *a* and *u* to *c*. Consider *y*^′^ edits occurring up to *d*_*u*_. Then the probability of having *x* total shared out-group edits is

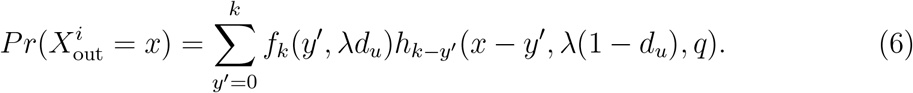

Now to work out 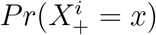, we consider all edits which arise on the path from *u* to *v*, as well as any additional shared sequence of edits which arises independently from *v* to *a* and *v* to *b*. We get

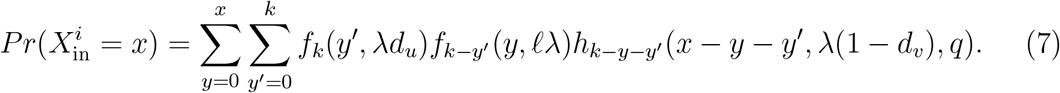

This is the probability that there were *y*^′^ ∈ [0, *k*] edits up to *d*_*u*_, *y* ∈ [0, *x*] edits along the path from *u* to *v*, and then *x* − *y* − *y*^′^ shared edits arising independently–and in sequence–along two paths of length 1 − *d*_*v*_.

The probability that the triplet is not resolvable is

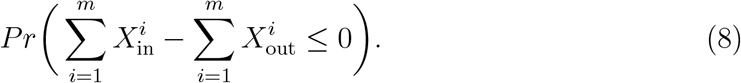

For small values of *mk*, we can compute this probability directly as an *m*-fold convolution. However, in general an approximation is necessary. Let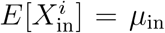, and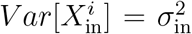, and similarly for the out-group. We then approximate Equation (8) by a normal distribution:

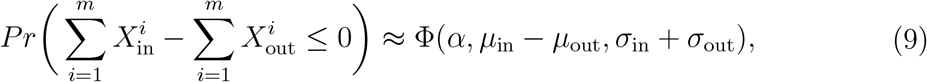

where ϕ denotes the cumulative density function of the normal distribution. We will take *α* = 1 for a conservative continuity correction, since taking *α* = 0.5 occasionally leads an overestimate of the reconstruction probability for small values of *m*. We now need to maximize the probability of not reconstructing the triplet over the entire tree. Let

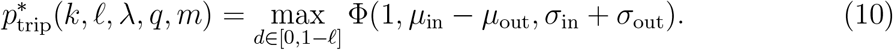

It is often the case that, due to the irreversible nature of edits and limited number of target sites, this minimum occurs when *d* is maximal. However, especially with higher values of *k, q* and lower editing rate *λ*, this minimum can occur higher in the tree. We now state our second result:

##### Result 2

*(B*_*q*_*) Given the tape mutation process described in Section 2, the probability of exactly reconstructing the phylogeny 𝒯 on n tips is at least*

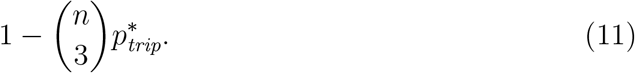

*where* 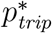*is the maximum probability of not resolving a triplet in 𝒯, and is approximated by Equation 10*.

Here, we have used Boole’s inequality to bound the probability that all triplets are resolved. This gives us an update to the bound provided in Equation (4) for the general case where 0 *< q <* 1. A significant limitation of using a normal approximation is that *m* needs to be large, although we can compute the exact probability in case of *m, k* small. In practice, over the parameter ranges we investigate, we find this approximation to be valid in terms of never overestimating the reconstruction probability. Further validations are given in Supplementary Figures B.10 and B.11.

### 3.2. Validation & Optimal Rates

We now compare our theoretical bounds to each other and to simulated data, and find explicit parameter regimes where exact reconstruction is likely (up to a given level of accuracy). For each experiment, on a fixed tree, we simulate tapes under the given parameters. Our goal is to assess the accuracy of trees inferred from these simulated tapes. Additional details of our simulation procedures are given in Appendix A.

First, we explore the accuracy of trees inferred using UPGMA to our theoretical bounds. To compute the simulated accuracy, we count the proportion of inferred trees which have the exact same topology as the true tree (i.e., a Robinson-Foulds distance of 0). Figure 2 illustrates our simulation results on a small tree (128 tips) under synchronous division. When looking at the distance between the reconstructed and true trees, we see that, as expected, there is a regime where the editing rate *λ* is too low, leading to a lack of information and failure to fully resolve the tree. Similarly, as *λ* is increased, there is again a loss of information as the tapes become pre-maturely saturated. The optimal editing rates occur at intermediate values, and the range of rates over which exact reconstruction occurs increases significantly as the number of target sites *k* per tape is increased.

**Figure 2.**
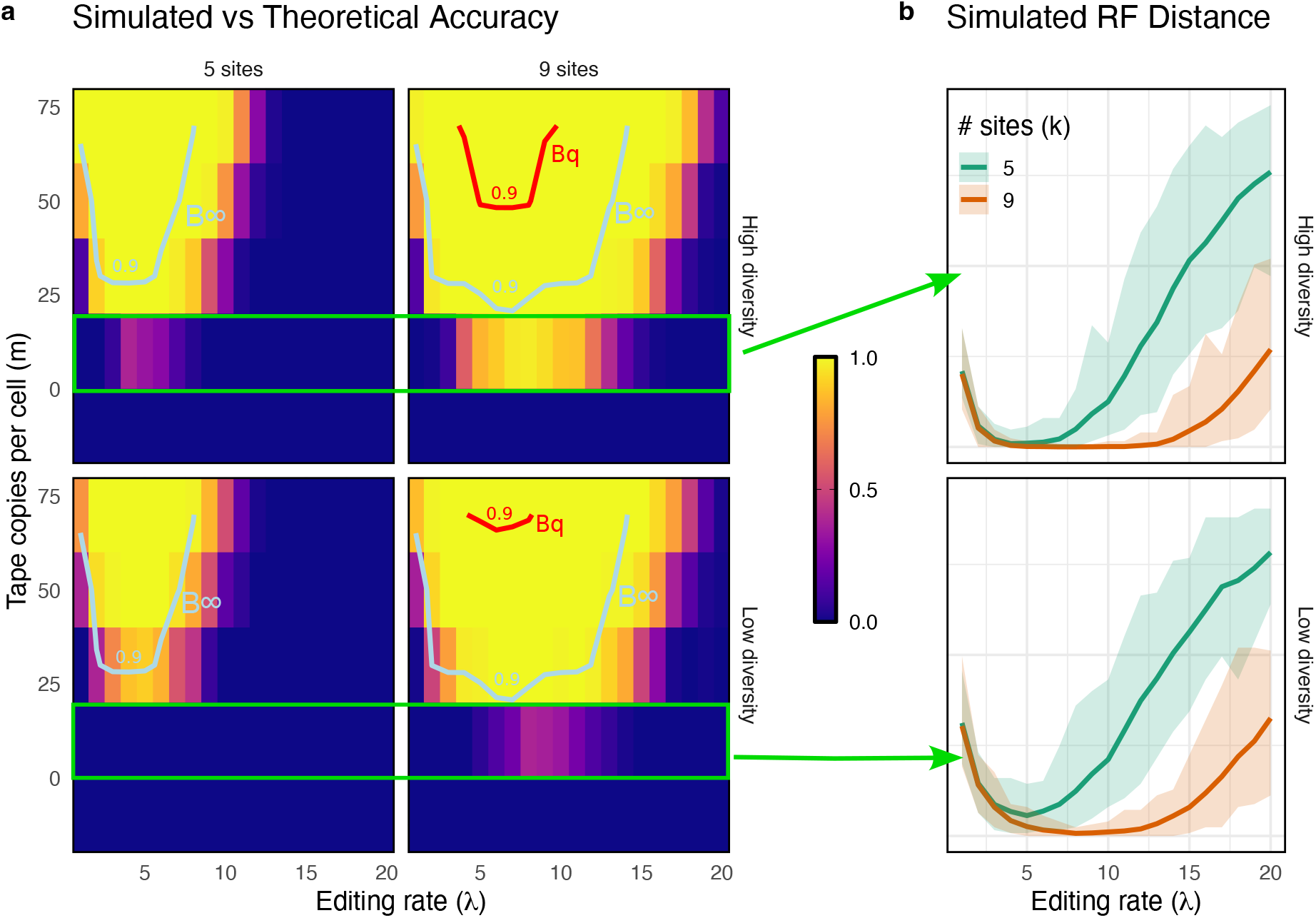
Simulated and theoretical probability of exact tree reconstruction. Phylogenies simulated under approximately synchronous division with *n* = 2^7^ tips. (a) Predicted 0.9 reconstruction accuracy based on Eqs. (4) and (11) (*B*_∞_ and *B*_*q*_, respectively). Heatmaps show the simulated accuracy (the proportion of exact reconstructions) over 100 simulations. Results are shown under low diversity (4 unique characters, *q* = 1*/*4) and high diversity (64 characters, *q* = 1*/*64) regimes. (b) Normalised Robinson-Foulds distance between UPGMA reconstruction and the true tree over 100 simulations of the mutation process (the tree remains fixed). The cases shown are with *m* = 10 tapes per cell (corresponding to the highlighted rectangular regions in panel (a)). Lines show median distance, ribbons show min/max distance.

We see that these general trends are reproduced by our theoretical estimates of accuracy (Figure 2a). Here we show the theoretical estimates for 90% accuracy given by Eqs. (4) and (11), as compared to the observed accuracy of the reconstructed trees over 100 simulations. We see that both *B*_∞_ and *B*_*q*_ predict optimal rates for intermediate values of *λ*, and that the range of optimal rates increases significantly with *k*, and also with the tape copy number *m*.

To further validate the bounds, we compare both bounds to a simulated triplet score over a wide range of parameter regimes. This score is computed as the proportion of simulated distance matrices for which all true triplets are resolvable. Our results are summarized in Figure 3 on fully sampled trees and also in Figure 4 for sampled trees. From Figure 3, we see that, in the high diversity regime, *B*_∞_ provides reasonable estimates for the expected accuracy, only slightly overestimating the accuracy under low rates as compared to the simulated triplet score. By contrast, *B*_*q*_ is much more robust, and we do not find any parameter regimes in which this bound is violated. It is, however, not particularly tight, and this is a problem which worsens with respect to the tree size.

**Figure 3.**
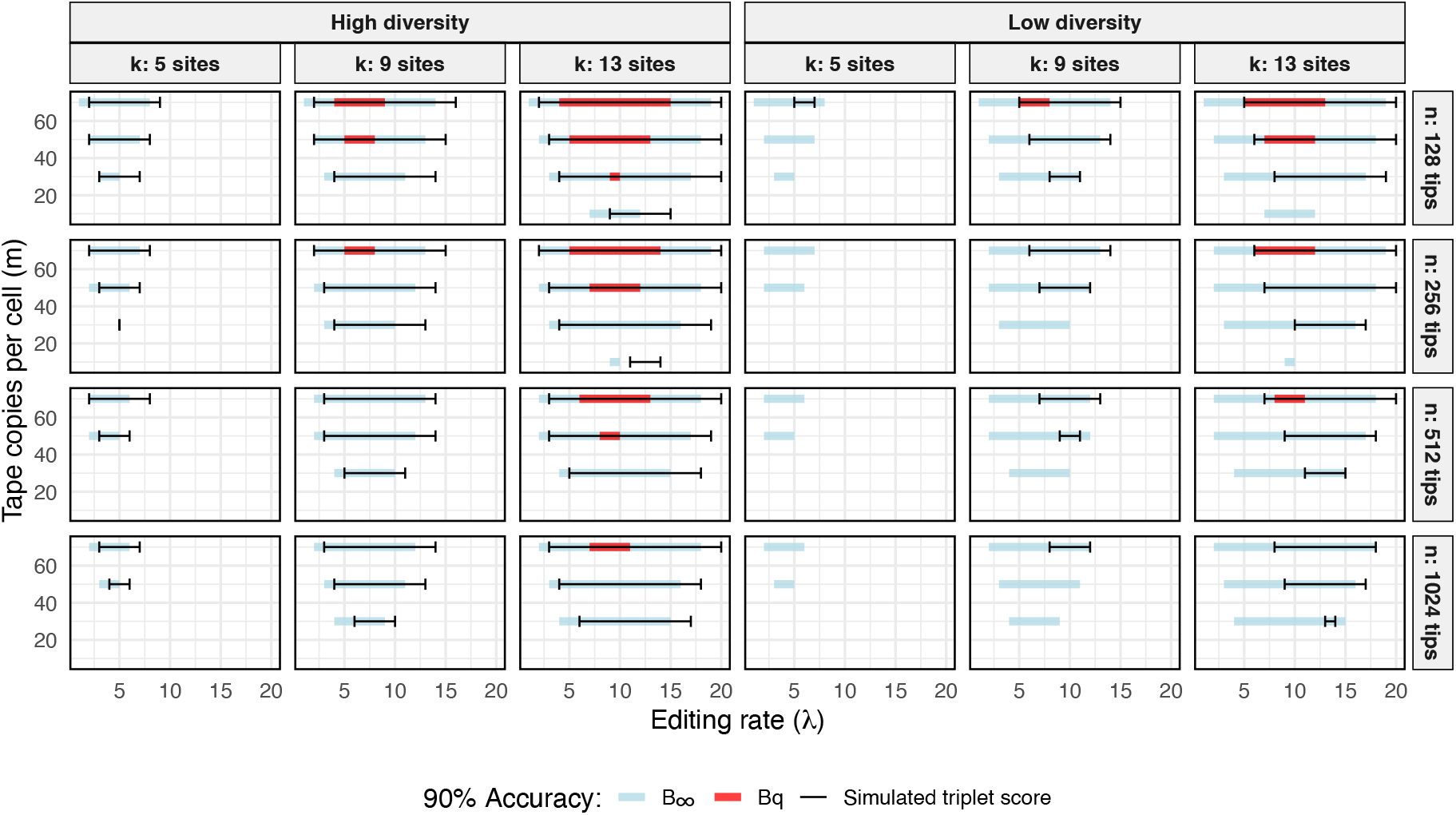
Simulated and theoretical parameter regimes for 90% reconstruction accuracy of the entire phylogeny under synchronous division. Editing rate *λ* is varied from 1–20. *B*_∞_ is given by Equation (4) and *B*_*q*_ by (11). Simulated accuracy is calculated as the proportion of simulations out of 100 where the triplet score is exactly 0 (i.e. all distances in the simulated distance matrix obey the true relationship). Results are shown under both low diversity (4 unique characters with *q* = 1*/*4) and high diversity (64 characters with *q* = 1*/*64) regimes. To compute *B*_∞_ and *B*_*q*_, we use the exact value of *ℓ* for each simulated tree.

**Figure 4.**
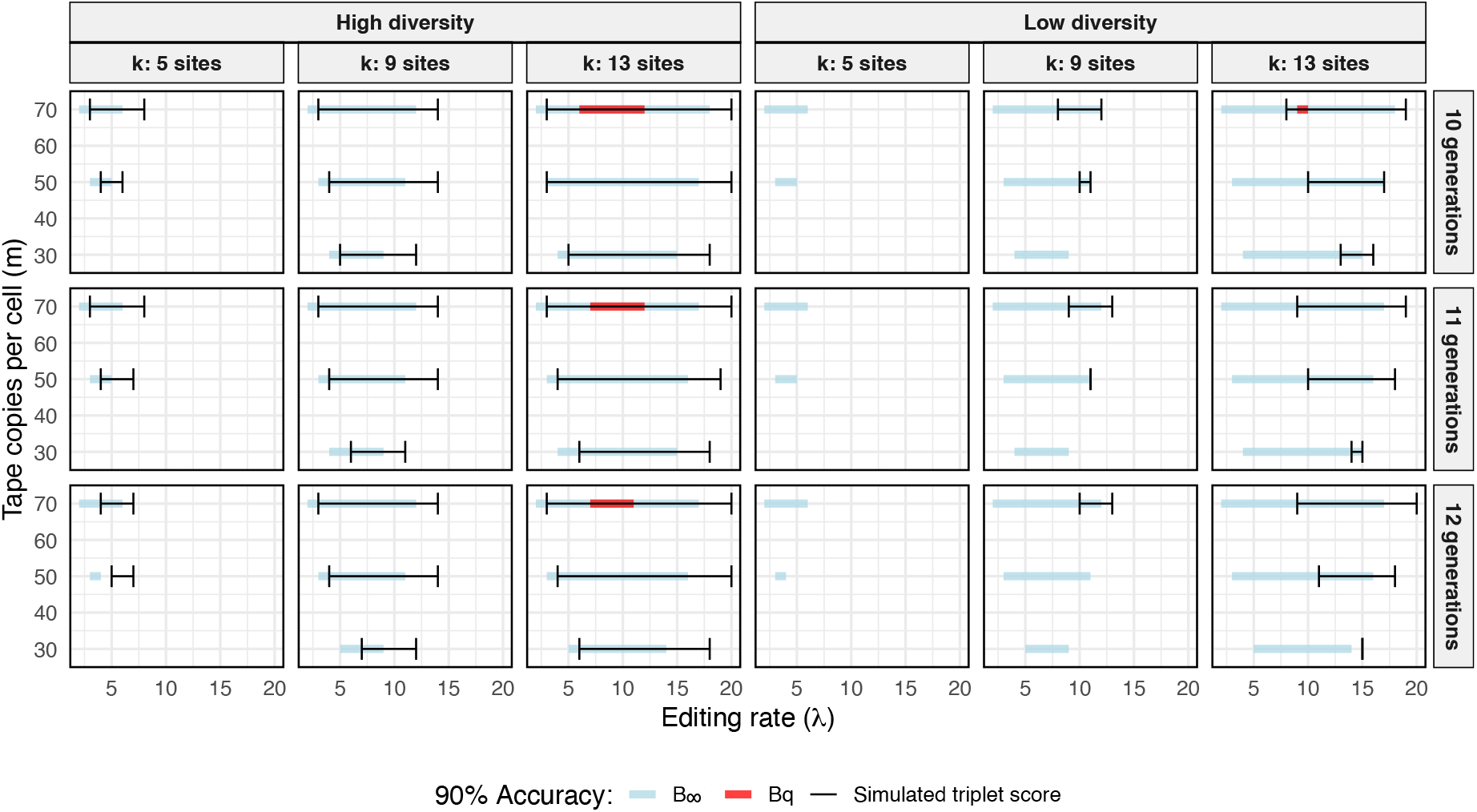
Simulated vs theoretical parameter regimes for 90% reconstruction accuracy on subsampled trees. Editing rate *λ* is varied from 1–20. Trees are generated for *n*_*g*_ generations under a Weibull branching process, and then randomly down sampled to approximately 100 tips. Theoretical reconstruction accuracy is determined by using Equations (4) and (11) for *B*_∞_ and *B*_*q*_ respectively, setting *n* = 100 and *ℓ* = 1*/*(*n*_*g*_ + 1). Simulated accuracy is calculated as the proportion of simulations out of 100 where the triplet score is exactly 0.

Both bounds become less tight in the sub-sampled case (Figure 4). In these experiments, we assume that the minimum branch length is the same as for the full process even though sampling may, in practice, skew the branch lengths longer. These results may then be more reflective of reality since, in practice, researchers may have an understanding of the minimum time to cell division, but not of the minimum branch length of a particular subsample of cells; in general, we cannot ignore the worst-case scenario.

We note again that *B*_∞_, which is only expected to be accurate as *q*→ 0, nonetheless appears to be a good approximation to the reconstruction accuracy as long as *q* is not too high. For high *q*, we see that *B*_∞_ consistently overestimates the reconstruction accuracy. We include *B*_∞_ in our results since it is a tight bound in the case of *q* = 0, whereas *B*_*q*_ is not. Therefore, estimates derived from *B*_∞_ may be useful in high diversity settings. By contrast, *B*_*q*_ provides a more robust bound on the reconstruction accuracy with respect to *q*.

#### 3.2.1. Scoring & Reconstruction methods

In Figures 3 and 4, we validated our bounds against a simulated “triplet score”, as this scoring method corresponds directly to our theory. To compute the triplet score, we simulate the tape evolution process on the true tree, then calculate the “simulated” distance matrix. We then check if all triplets are resolvable (Eq. (**??**)) based on the simulated distances.

Another way to assess the accuracy is to reconstruct a tree based on the simulated distances (for example, what was done in Figure 2). To get a measure of accuracy that corresponds with our theory, we look at the proportion of inferred trees which have the exact same topology as the true tree. That is, where the Robinson-Foulds (RF) distance is exactly 0. Theoretically, a tree reconstruction method based on triplets, such as a naïve triplet method (Wang et al., 2023), would provide the same results as the triplet score.

In practice, we expect any appropriate tree-reconstruction method to perform better than a triplet-based method. For example, in Figure 5 we compare the triplet score to scores based on the commonly-used UPGMA algorithm and also the NeighbourJoining method. We see that, as expected, both tree-based scores are more optimistic than the triplet scores. We would expect a likelihood-based approach (such as Seidel et al. (2024)) to perform even better than these two distance-based approaches, although a detailed comparison of tree-reconstruction methods for these data is left for future work. In particular, here, we are only interested in the tree topology, and not on more detailed measures of tree reconstruction (including branch lengths etc.).

**Figure 5.**
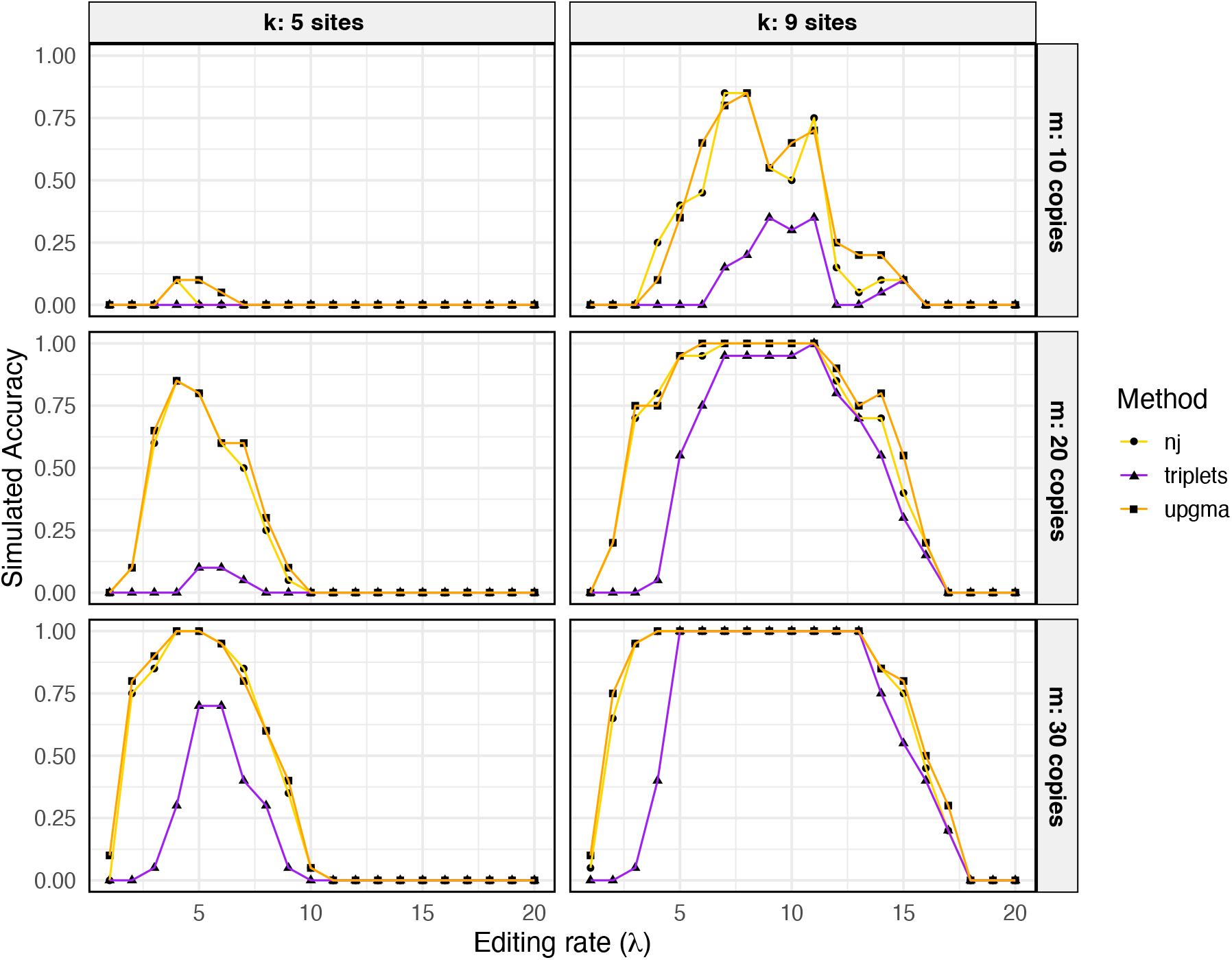
Comparison of triplet score to scores from two different tree reconstruction methods. Triplet score computes the accuracy as the proportion of simulations where all triplets can be resolved exactly. For the neighbour-joining(nj) and UPGMA-based scores, the score is computed as the proportion of simulations in which the inferred tree topology is correct (i.e. a RF distance of 0 to the true tree). Experiments are repeated 20 times on a tree of 128 tips with *ℓ* ≈ 1*/*9.

### 3.3. Minimum number of copies

We now investigate the minimum copy number *m* required to generate data with a given reconstruction accuracy. Even though increasing the number of target sites per tape *k* will always lead to the most drastic improvements, in practice this parameter is difficult to increase (Liao et al. (2024)). Therefore, we instead consider how many copies of each tape we need per cell to achieve a given accuracy according to our theoretical bounds, constrained to given values of *k* and *ℓ* (the desired tree resolution). We further assume that the editing rate *λ* can be tuned as needed.

We illustrate how to use Equations (4) and (11) to get an estimate for the minimum number of tapes *m*^∗^ required to reach a desired level of accuracy *ϵ*. Our approach is to first use Eq. (3) to find the optimal editing rate (note that this does not depend on *m*), and then to use this rate in both Eqs.(4) and (11) (*B*_∞_ and *B*_*q*_ respectively) to find *m*^∗^.

We first find *λ*^∗^ by maximising Eq. (4) over the entire tree (*d* = 1 − *ℓ*),

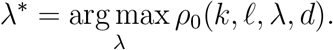

Then the estimate for *m*^∗^ using *B*_∞_ is *m*^∗^ = ⌈*m*⌉ where

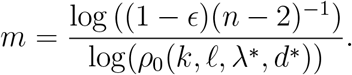

The estimate for *m*^∗^ using *B*_*q*_ is found numerically by finding where Eq. (11) is equal to *ϵ*, using the previously computed value for *λ*^∗^. Note that this includes an extra optimization step to solve Eq. (10). Based on the results in Figures 2 and 3, we expect *B*_*q*_ to overestimate the minimum required copy number, while *B*_∞_ can provide a more realistic estimate but may underestimate this value if *q* is not small enough, and especially for lower *λ* values.

These results are shown over a range of experimental parameters *k, ℓ, n* in Figure 6. We interpret *ℓ* here as the minimum desired resolution rather than the shortest possible branch length, and note that these results are much more sensitive to *ℓ* rather than *n*. Even though it is not robust with respect to *q, B*_∞_ can be used to get a realistic estimate of *m*^∗^. For example, if a given system has *k* = 5 sites, then at least ca. 30 tapes would be required to accurately resolve a fully sampled phylogeny over 9 generations of uniform cell divisions (*ℓ* ≈0.1, *n* = 512), or at least 24 if only 100 cells are sampled. If we decrease *ℓ* to 0.05, then at least 50 copies would be needed to achieve the same level of accuracy over 100 cells. We have assumed here that the exact rate *λ*^∗^ is achievable, and so these are optimistic values. We might furthermore expect Eq. (11) to have different optimal value *λ*^∗^, but as long as *q* is not too high we expect these values to be close. These results also highlight the non-tightness of *B*_*q*_, as it does not reduce to *B*_∞_ when *q* = 0, due to the fact that we must bound over all triplets when computing *B*_*q*_.

**Figure 6.**
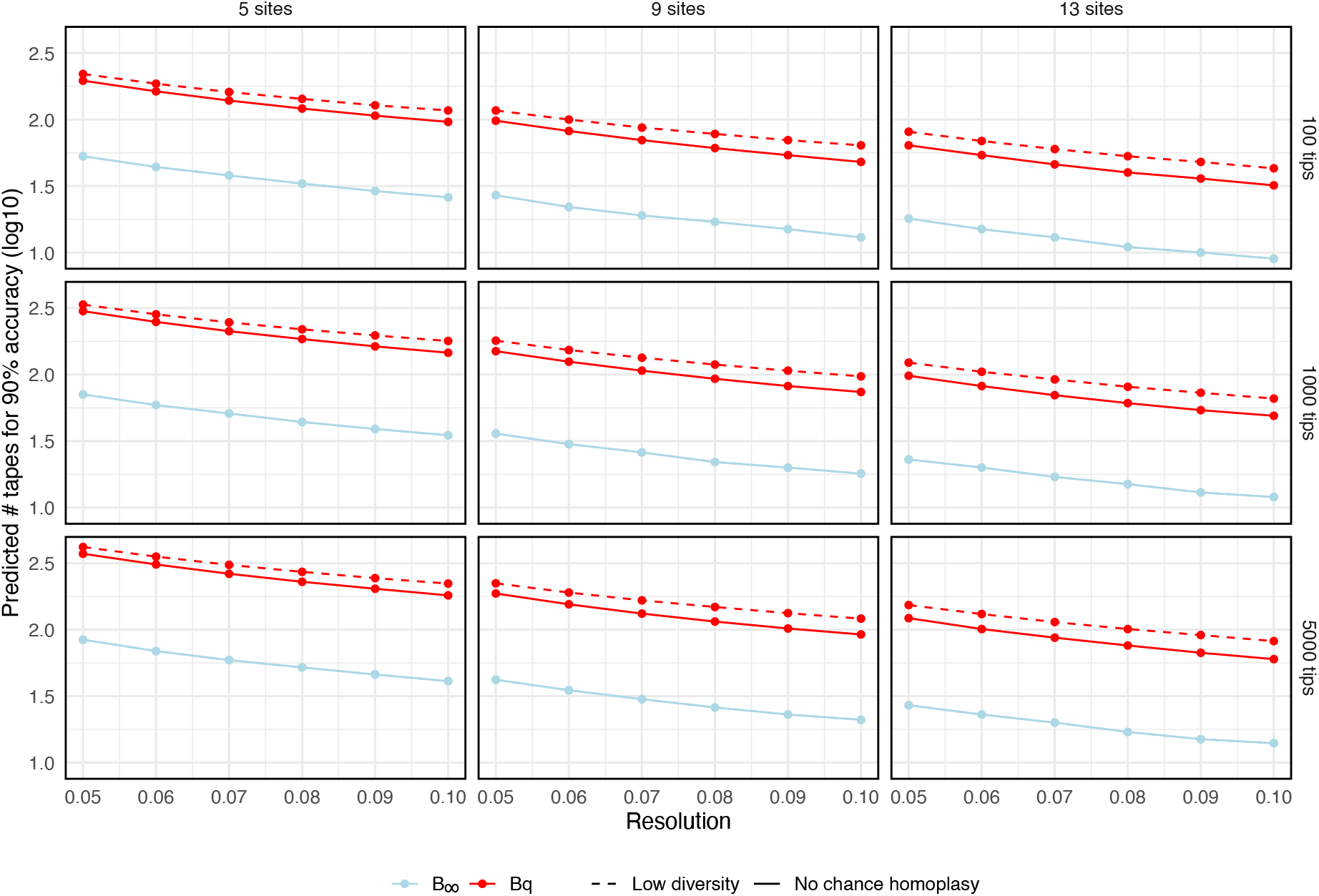
Predicted value for *m*^*∗*^, the minimum copy number required for 90% accuracy. The parameter *n* is the number of sampled tips, *k* is the number of target sites per copy, and the resolution corresponds to parameter *ℓ*. Low diversity corresponds to *q* = 0.25, while no homoplasy corresponds to *q* = 0.

### 3.4. Multiple editing rates

This analysis further provides some insights into when multiple editing rates may help resolve the phylogeny. In particular we investigate the case where early development is characterised by short cell cycles, as this leads to a case where the phylogeny is more difficult to resolve (shorter branches deep in the tree). Consider a population of cells which initially divide at some rate *αr* for *n*_1_ generations, and then at rate *r* for the rest of the experiment. We consider only the case where *α <* 1, since this is the scenario where it would be most beneficial to have multiple editing rates to resolve both stages of growth. We assume that in both stages the cells divide approximately synchronously. We will assume that *q* is small enough that we an use Eq. (4) to provide a realistic estimate for the accuracy of the reconstruction, although we could perform a similar analysis using Eq. (11). Given *k, m, ℓ*_1_, *ℓ*_2_, *n*_1_, *n*_2_ our goal is to find rates *λ*_1_, *λ*_2_ and copy numbers *m*_1_, *m*_2_ so as to achieve a predetermined reconstruction accuracy, *ϵ*.

First, we compute the probability of reconstructing the entire phylogeny, with all *m* tapes and minimum branch length *ℓ* = *ℓ*_1_. If this yields the desired accuracy, we do not need multiple editing rates. If it does not, we take the following approach, following a similar procedure to that outlined in Section 3.3. Let

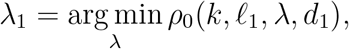

where *ρ*_0_ is given by Eq. (3), and *d*_1_ = (*n*_1_ − 1)*ℓ*_1_. Next, let *m*_1_ = max(*m*, ⌈*m*^∗^⌉) where

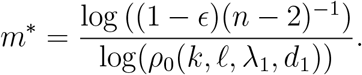

If *m*_1_ *< m*, let *m*_2_ = *m* − *m*_1_ and similarly let

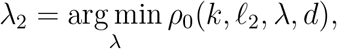

where now *d* = 1- *ℓ*_2_. In this way, we get two rates with *λ*_1_ *> λ*_2_, and prioritise resolving early stage divisions. Note however that this procedure does not have a guarantee on the accuracy of the entire tree reconstruction.

Figure 7 shows how this approach can resolve the phylogeny in scenarios where doing so would be impossible with just a single rate. We compare the results of experiments using two rates (as described above) to experiments which use only a single rate. In the single rate case, we optimise Eq. (3) fixing the minimum branch length to be *ℓ*_1_. We see that using two rates can lead to significantly better final accuracy over using just a single rate. As expected, this approach becomes more beneficial as *α* decreases; although, as *ℓ*_1_ decreases, it also becomes more difficult to resolve the phylogeny. This is especially true as we increase the duration of the fast growth stage. We note that here we keep the total number of tapes fixed, and we have no constraints on the editing rate(s).

**Figure 7.**
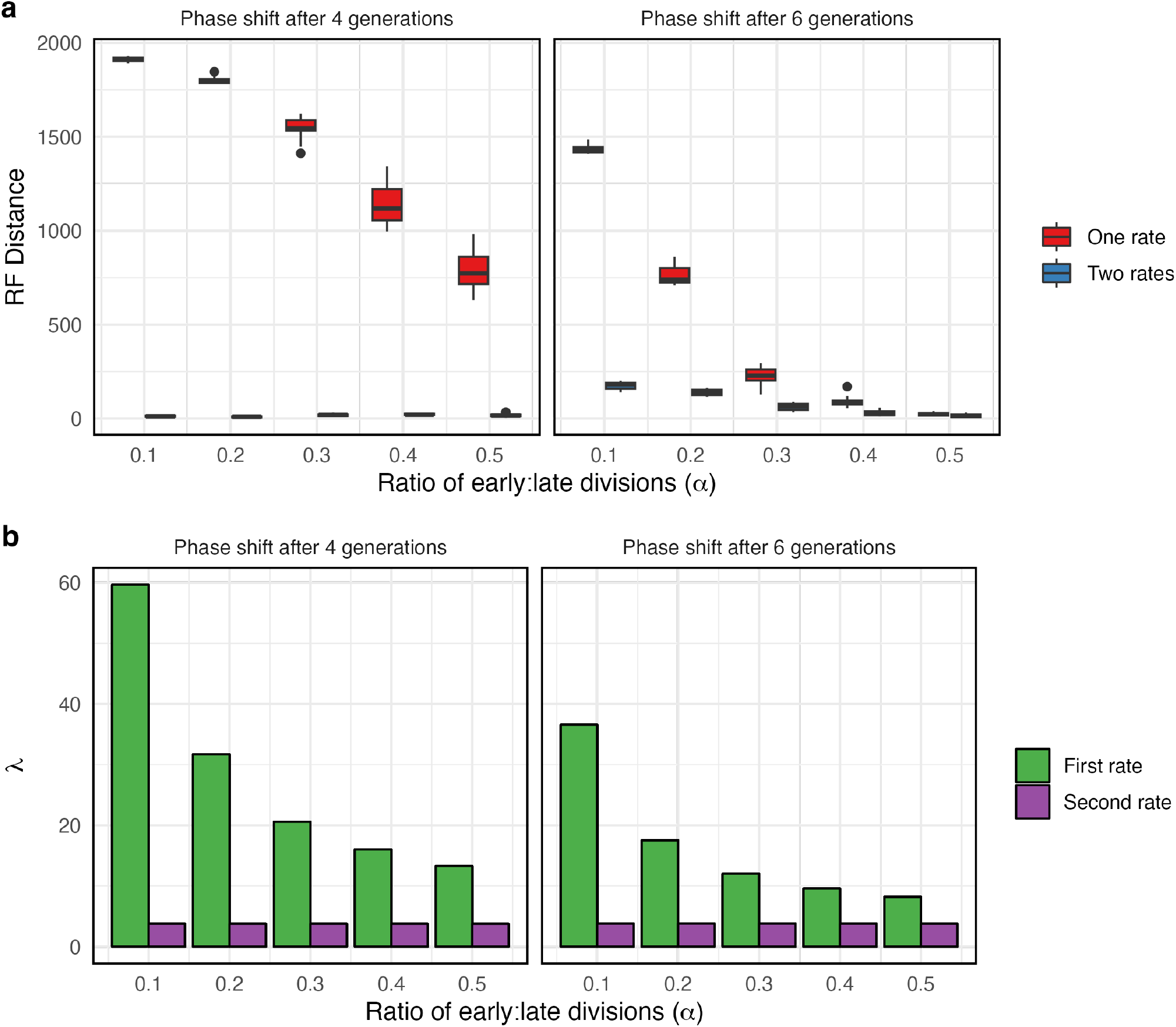
Multiple rates can be beneficial for resolving fast, early stage divisions. (a) Improved accuracy by using two rates, following the procedure outlined in Section 3.4. Fixed parameters: *k* = 5, *ℓ*_1_ = *αℓ*_2_, *q* = 1*/*64, *m* = 20, *n* = 1024. Tree is simulated under approximately uniform division, with 10 generations total and phase shift occurring after either 4 or 6 generations. Results show the Robinson-Foulds distance between the true tree and the UPGMA reconstructed tree over 100 simulations. (b) The difference in magnitude between the two rates. Where only a single rate is used, the larger rate is chosen.

### 3.5. Insertion probabilities

Our theoretical result suggest that the theoretical reconstruction accuracy does not depend on the exact distribution of insertion probabilities, only on the so-called collision probability *q*. This is readily verified through simulations. In Figure 8, we compare two experimental set-ups with identical values of *q*, but one with a skewed insertion distribution, and another with a uniform insertion distribution. As expected, we see no significant differences between the two over 50 simulations.

**Figure 8.**
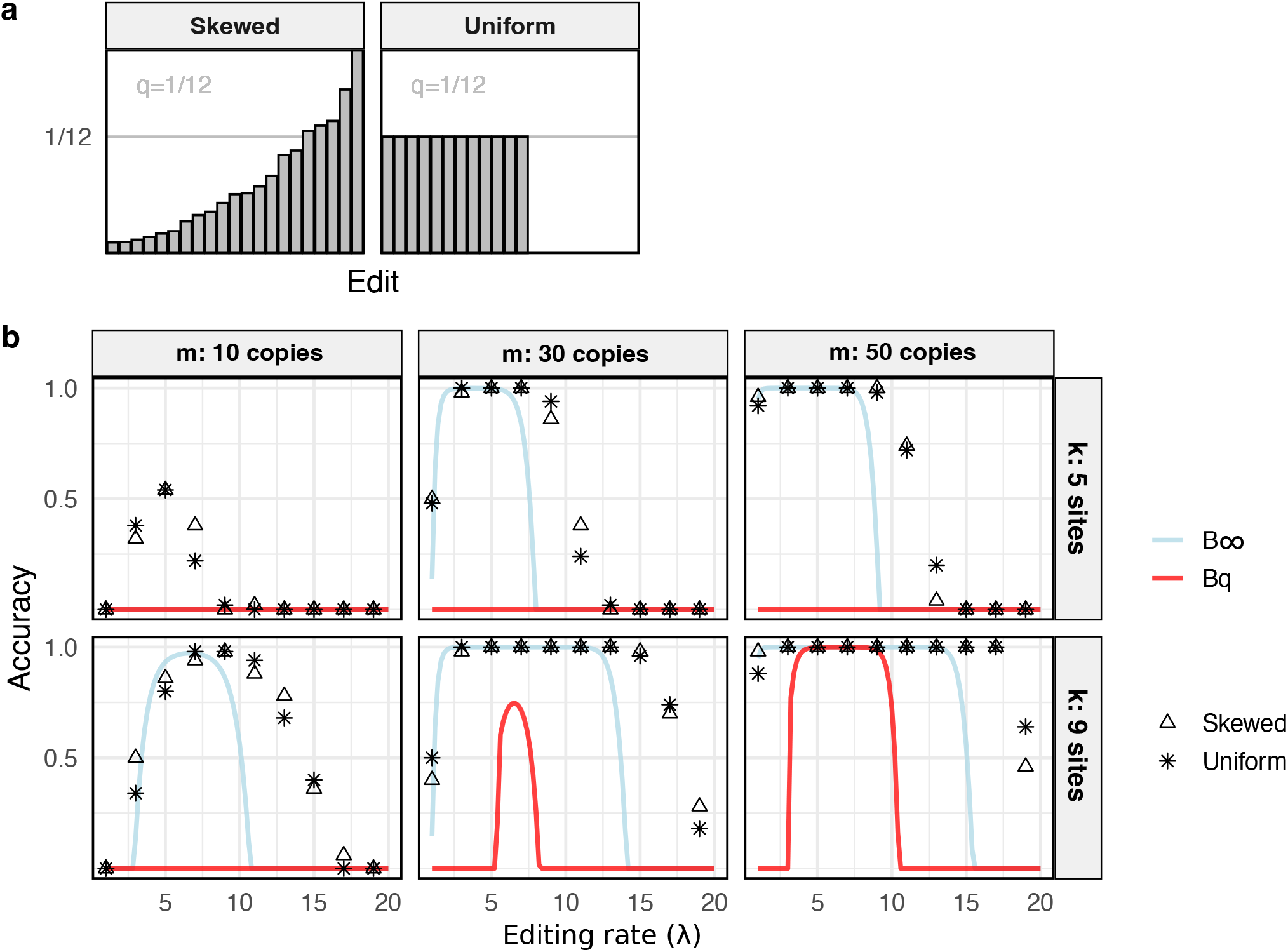
Accuracy does not depend on the distribution of edits. (a) Two distributions of insertion probabilities, each with *q* = 1*/*12: On the left, a skewed distribution with 21 characters, and on the right, a uniform distribution on 12 characters. (b) Results over 50 simulations on a single tree. Simulations are identical up to the edit distributions. Tree has 2^6^ tips with *ℓ* ≈1*/*7. Accuracy is calculated as the proportion of inferred UPGMA trees which have the exact true topology (RF distance 0).

One potential concern when applying Equation (11) or (3) in practice is the requirement of prior knowledge of certain parameters, including *q* (the probability that two independent editing events insert the same character). This is a property of the insertion rate distribution of the editing system. If *j* is the number of unique possible characters, then 1*/j* ≤ *q <* 1. We show how *q* can be readily estimated from the barcode “residuals”: consider two tapes, *s*_1_ and *s*_2_, of length *k*. Then remove the sequentially shared edits, together with the first non-shared edit. Then *q* can be estimated from the proportion of remaining edits which are shared across all barcodes in the experiment (Figure 9).

**Figure 9.**
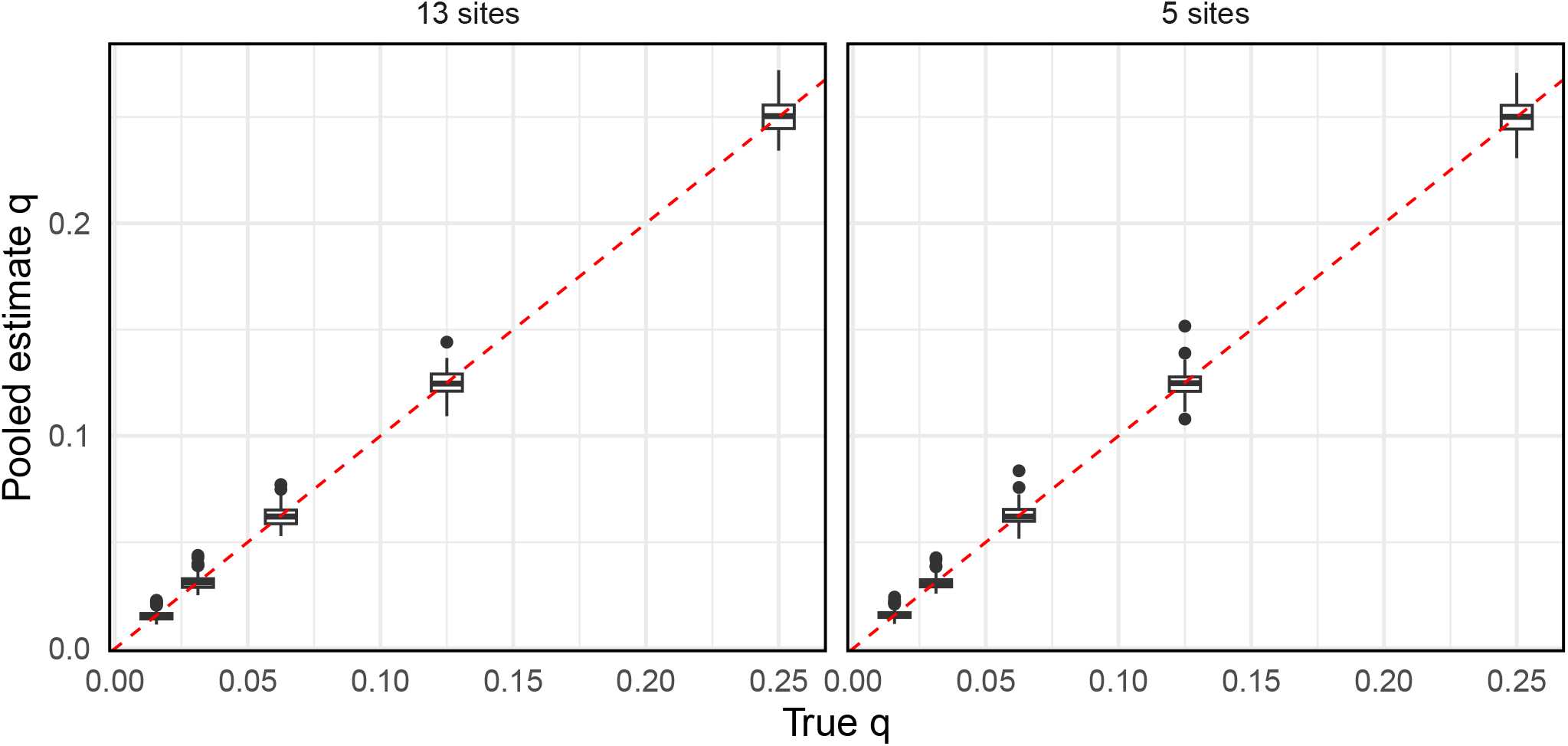
Estimated vs true *q* over 100 simulations, for *k* = 13 (left) and *k* = 5 (right). Here, *m* is chosen so that a given set-up can achieve approx. 90% accuracy.

## 4. Discussion

Emerging technologies such as DNA Typewriter (Choi et al., 2022) and PeChyron (Loveless et al., 2021) have significant potential to generate data which can be used to accurately reconstruct cellular phylogenies. We have explored, through simulation and theory, when sequentially edited systems lead to accurate reconstruction, and have provided examples of how our theoretical results could be used to inform experimental design. We furthermore shed light on regimes wherein computationally efficient, distance-based reconstruction methods can be used to accurately reconstruct the cellular phylogeny. Our model assumes that the editing process is irreversible, in line with other models for CRISPR recording devices, and also sequential. We generally assume that there is a finite space of possible edits and that there are a limited number of target sites, although both of these parameters could be made arbitrarily large. We furthermore consider the case where many tapes can be inserted into a cell’s genome. Therefore, our model is appropriate for a recorder system that is sequential, highly multiplexible and robust. This work expands upon previous research such as Wang et al. (2023) and Salvador-Martínez et al. (2019) which investigate the information content in data from earlier generation CRISPR recorders.

A limitation of our approach is that we do not explicitly model sources of error. In protocols such as Typewriter, the most common sources of sequencing error–single base-pair substitutions–can be estimated through simple post-processing steps (Liao et al., 2024), and so in this work we do not explicitly model this random source of error. Earlier molecular recording systems additionally suffered from data loss and corruption in the form of double-resection events (Salvador-Martínez et al., 2019; Feng et al., 2021). These events occur when two recording sites are cut simultaneously, creating a deletion that spans the intermediate sites, and erasing any lineage information that may have been present at those sites. Both Typewriter (Choi et al., 2022; Liao et al., 2024) and PeChyron (Loveless et al., 2021) evade this source of missingness through the use of prime editor, which introduces only small, single-stranded nicks into the genome. Accordingly, we do not model this source of missing data. However, it is possible for an inserted recorder to become silenced or unexpressed over time (Cabrera et al., 2022; Chen et al., 2022; Choi et al., 2022). We assume that these occurrences do not depend on nor influence other model parameters, and so we do not explicitly model this process here. Our parameters *m* and *n*, the number of required tape copies recovered from *n* sequenced cells, are therefore idealised and in practice would correspond to a subset of the initial 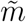number of inserted tapes and final number *ñ* of cells. Then 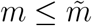would correspond to the number of tapes which are recovered in all *n* ≤ *ñ* cells.

Another assumption of our model is that the editing rate for a given tape is constant over the entire duration of the experiment. In reality, there are a number of factors which could cause this rate to be non-constant. In some recorder systems, it is known that the editing rate decreases over time due to decay of the editing enzymes. However, this is not expected to be a particular concern in systems such as Typewriter where cells are engineered to express the editing enzyme (Liao et al., 2024). On the other hand, this setup means that recording cannot be induced at a later stage of cell development. Site-of-integration effects could, however, change the effective editing rate (and also make the tape difficult to recover (Choi et al., 2022)). While we address this issue somewhat by displaying the range of rates over which accurate reconstruction can be achieved, the constant-rate assumption could be addressed directly in future work.

Our results can be used to assess whether or not current experimental technologies are close to being able to achieve high reconstruction accuracy or not. As of recent publications (Choi et al., 2022; Liao et al., 2024), the Typewriter protocol is able to stably integrate tapes of approx. 5 target sites with 64 possible edits and up to approximately 20 copies. In our notation, this corresponds to *k* = 5, *j* = 64 and *m* = 20. We note that the distribution of edits may be skewed in practice, and so it is likely that *q* is greater than 1*/*64. Our results indicate that such experiments are generally not yet in a regime where exact reconstruction is possible for trees on the order of 1000 cells (Figure 6), but that accurate reconstruction may be tangible as long as cell division rates are not too high. For example, in an optimistic setting of 10 synchronous generations (corresponding to *ℓ* ≈ 0.09), Figure 6 predicts that approximately 60 tapes are needed to guarantee a reconstruction accuracy of at least 90% (according to *B*_∞_). We furthermore note that it may not always be practical to finely tune parameters such as the editing rate (Liao et al., 2024). How well this editing rate can be tuned would have implications for determining sufficient values for *k* and *m*. Given that the information content of these data increase greatly as the number of target sites is increased, the ability of PeChyron to generate new target sites during the recording process appears to be highly promising (Loveless et al., 2021), although we note that the current information capacity of this system is similar to that discussed above; while Typewriter is most limited in the number of target sites, PeChyron is limited in its ability to tune the editing/cut rate (and thereafter create a new target site).

We see a variety of avenues for future work and alternative approaches. Such work could, for example, address the robustness of *B*_*q*_ with respect to the Normal approximation, and/or find a better approximation for the distribution in Eq. (8). Alternatively, it may also be possible to determine a bound which reduces to *B*_∞_ as *q* → 0 (unlike our *B*_*q*_ here) but is still valid when the chances of homoplasy are not negligible (which *B*_∞_ assumes). In our current approach, our bounds are always based on the worst-case scenario, which is inevitable since we are concerned with exact reconstruction. However, a probabilistic approach to a relaxed problem could be an avenue for future work. An alternative approach could take into account both the underlying cellular dynamics and the effect of sampling on the branch length distribution.

We have illustrated here, through both simulations and theory, the information capacity of sequential lineage recorders, and our results suggest that the goal of highly accurate reconstruction in this setting is feasible. In particular, while our results indicate that these technologies are not yet able to fully resolve phylogenies on the order of thousands of cells, we provide examples of how our theoretical results can be used to help guide experiments and improve the accuracy of the resulting phylogeny.

## Acknowledgments

TS received funding from the European Research Council (ERC) under the European Union’s Horizon 2020 research and innovation programme grant agreement No 101001077 (PhyCogy). The authors would like to thank Antoine Zwaans, Marcus Overwater and Timothy G. Vaughan for helpful discussions and comments on the manuscript.

## Appendix A. Supplementary Methods

We simulate ultrametric cellular phylogenies with *n* tips using TreeSimGM (Hagen and Stadler, 2018). Birth (cell division) events occur according to a Weibull process with scale parameter *k*. When *k* = 1, this reduces to an exponential distribution, and as *k* grows large, division times become more and more synchronous. For “synchronous trees”, unless otherwise stated we take *k* = 200. We assume that cell death is negligible. Each phylogeny is scaled so that the depth of the tips is equal to 1. We then simulate an independent tape mutation process along this phylogeny using Ape (Paradis and Schliep, 2019), and repeat *m* times along the same phylogeny, recording the data from only at the tips of the tree. We then compute the resulting distance matrix and build the corresponding phylogeny using UPGMA. The reconstructed tree is said to be accurate if the Robinson-Foulds distance to the true tree is exactly 0. Alternatively, we can use a Triplet score which takes the simulated distance matrix and checks if every triplet is resolvable. In either case, we count the proportion of simulations which result in an exact reconstruction. In our simulations, we typically assume a uniform distribution over the space of possible edits, so that *q* = 1*/j*, although this need not be true in practice. The code required to run these simulations is available at: https://github.com/nmulberry/Strategies-Sequential-Editing.

## Appendix B. Supplementary Figures

**Figure B10.**
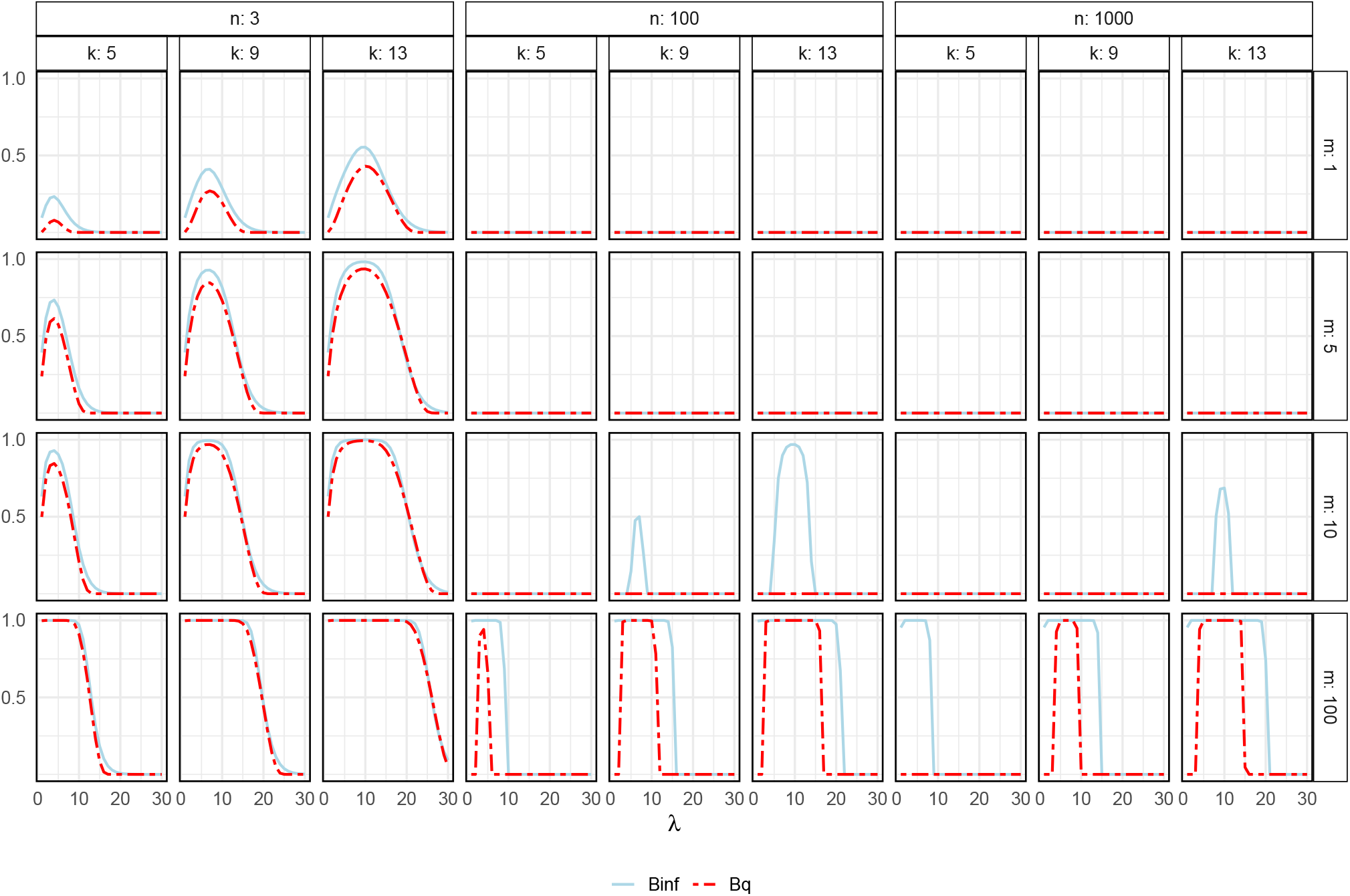
Validity of the approximation Eq. (9) as *m* increases with *q* = 0. We expect *B*_*q*_ to approximate *B*_∞_ as *m* → ∞ and for *n* = 3 (a single triplet). For *n >* 3, we see that *B*_*q*_ underestimates compared to *B*_∞_. We find no parameter regimes where *B*_*q*_ overestimates the reconstruction probability. Fixed *ℓ* = 0.01.

**Figure B11.**
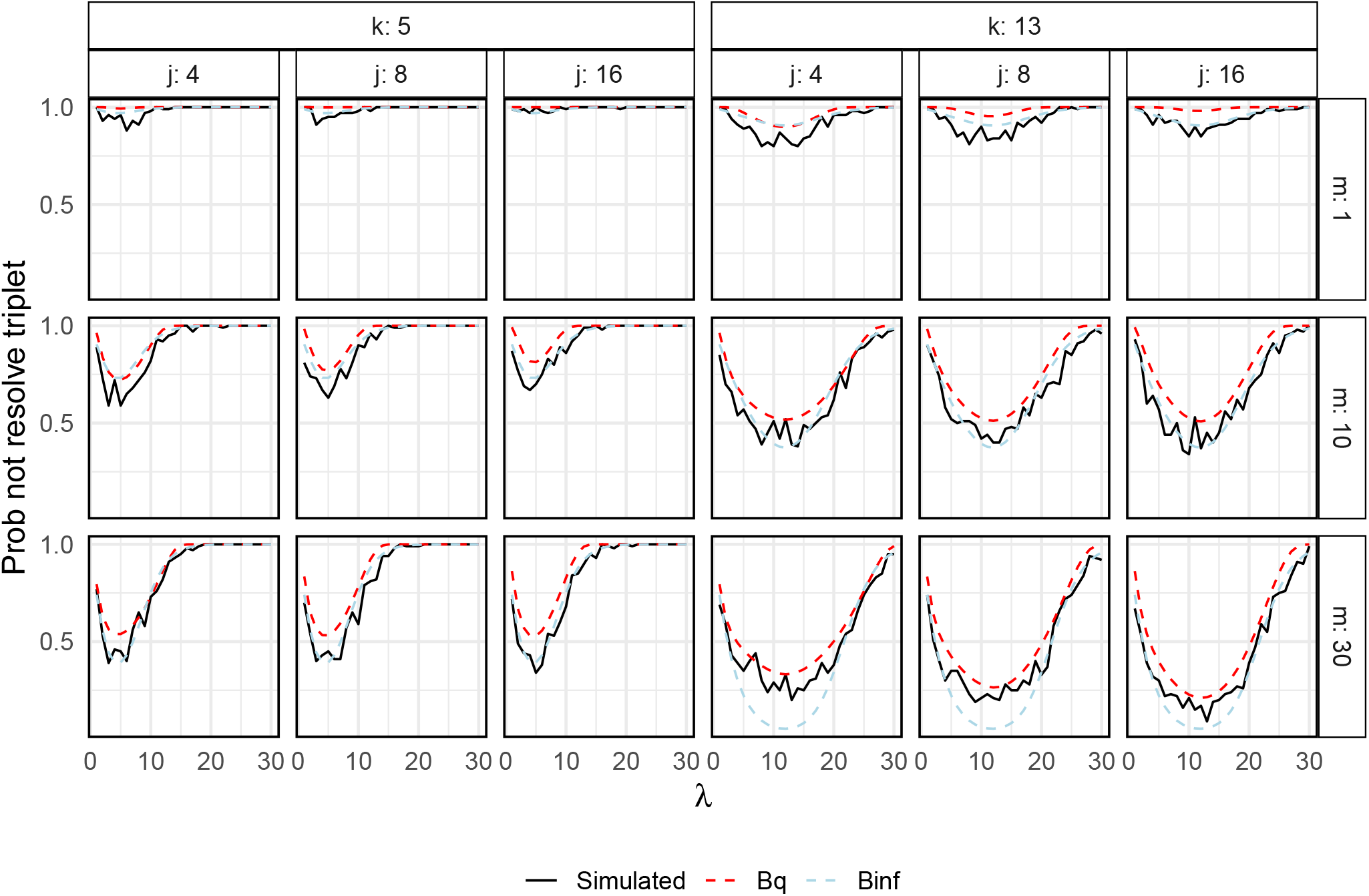
Probability of resolving a single triplet (*a, b*| *c*). Minimum branch length *ℓ* varied, fixed depth *d* = 0.8. We simulate *m* tapes of length *k*, starting at the root state **0**^*k*^. For each tape, from *t* = 0 to *t* = *d* = 0.8, we simulate a single sequence that will be shared between all three nodes. From *t* = 0.8 to *t* = 0.8 + *ℓ*, we allow two sequences to evolve independently, and then from *t* = 0.8 + *ℓ* to *t* = 1, we consider three sequences to evolve independently. For each scenario, we repeat over 100 simulations. We then count the proportion of simulations in which the triplet is resolved. This provides further validation of Eqs. (9) and (3). Fixed *ℓ* = 0.01.

## References

Tanja Stadler, Oliver G Pybus, and Michael PH Stumpf. Phylodynamics for cell biologists. Science, 371(6526):eaah6266, 2021.

Aaron McKenna and James A Gagnon. Recording development with single cell dynamic lineage tracing. Development, 146(12):dev169730, 2019.

Cheng Chen, Yuanxin Liao, and Guangdun Peng. Connecting past and present: single-cell lineage tracing. Protein & Cell, 13(11):790–807, 2022.

Junhong Choi, Wei Chen, Anna Minkina, Florence M Chardon, Chase C Suiter, Samuel G Regalado, Silvia Domcke, Nobuhiko Hamazaki, Choli Lee, Beth Martin, et al. A time-resolved, multi-symbol molecular recorder via sequential genome editing. Nature, 608(7921):98–107, 2022.

Theresa B Loveless, Courtney K Carlson, Catalina A Dentzel Helmy, Vincent J Hu, Sara K Ross, Matt C Demelo, Ali Murtaza, Guohao Liang, Michelle Ficht, Arushi Singhai, et al. Open-ended molecular recording of sequential cellular events into dna. bioRxiv, pages 2021–11, 2021.

Mareike Fischer and Mike Steel. Sequence length bounds for resolving a deep phylogenetic divergence. Journal of theoretical biology, 256(2):247–252, 2009.

Alex Dornburg, Zhuo Su, and Jeffrey P Townsend. Optimal rates for phylogenetic inference and experimental design in the era of genome-scale data sets. Systematic Biology, 68(1):145–156, 2019.

Jeffrey P Townsend and Francesc Lopez-Giraldez. Optimal selection of gene and ingroup taxon sampling for resolving phylogenetic relationships. Systematic Biology, 59(4):446–457, 2010.

Robert Wang, Richard Zhang, Alex Khodaverdian, and Nir Yosef. Theoretical guarantees for phylogeny inference from single-cell lineage tracing. Proceedings of the National Academy of Sciences, 120(12):e2203352120, 2023.

Irepan Salvador-Martínez, Marco Grillo, Michalis Averof, and Maximilian J Telford. Is it possible to reconstruct an accurate cell lineage using CRISPR recorders? Elife, 8:e40292, 2019.

Robert R Sokal and Charles D Michener. A statistical method for evaluating systematic relationships. 1958.

Sophie Seidel, Antoine Zwaans, Junhong Choi, Samuel Regalado, Jay Shendure, and Tanja Stadler. Leveraging sequential lineage recording data with SciPhy: A Bayesian approach for estimating single cell trees and dynamics. in preparation, 2024.

Hanna Liao, Junhong Choi, and Jay Shendure. Molecular recording using dna type-writer. Nature Protocols, pages 1–30, 2024.

Jean Feng, William S DeWitt III, Aaron McKenna, Noah Simon, Amy D Willis, and Frederick A Matsen IV. Estimation of cell lineage trees by maximum-likelihood phylogenetics. The annals of applied statistics, 15(1):343, 2021.

Alan Cabrera, Hailey I Edelstein, Fokion Glykofrydis, Kasey S Love, Sebastian Palacios, Josh Tycko, Meng Zhang, Sarah Lensch, Cara E Shields, Mark Livingston, et al. The sound of silence: Transgene silencing in mammalian cell engineering. Cell Systems, 13(12):950–973, 2022.

Oskar Hagen and Tanja Stadler. TreeSim GM: Simulating phylogenetic trees under general Bellman–Harris models with lineage-specific shifts of speciation and extinction in R. Methods in ecology and evolution, 9(3):754–760, 2018.

Emmanuel Paradis and Klaus Schliep. ape 5.0: an environment for modern phylogenetics and evolutionary analyses in R. Bioinformatics, 35(3):526–528, 2019.

